# The conservation status of Costa Rican velvet worms (Onychophora)

**DOI:** 10.1101/493833

**Authors:** Bernal Morera-Brenes, Julián Monge-Nájera, Paola Carrera Mora

## Abstract

Charismatic species, like the panda, play an important role in conservation, and velvet worms arguably are charismatic worms. Thanks to their extraordinary hunting mechanism, they have inspired from a female metal band in Japan, to origami worms in Russia and video game monsters in the USA.

**Objective:** To assess their conservation status in Costa Rica.

**Methods:** we located all collection records of the 29 known species from the Onychophora Database in the map of the Costa Rican Conservation Network.

**Results:** We found that seven species are protected inside forest reserves, five in Protected Zones, four in Wildlife Refuges, two in National Parks and one, *Principapillatus hitoyensis,* in a strictly pristine Biological Reserve. The largest species in the world, *Peripatus solorzanoi,* occurs both inside a Forest Reserve and in protected private land. Protection inside Costa Rican nature areas is enforced year round by personnel that includes armed guards, and is supported by educational programs in surrounding communities. Twelve species have not been found in protected areas, but in Costa Rica, all biological species, named and unnamed, are protected by law and cannot be legally collected, or exported, without technically issued permits.

**Conclusion:** Like in the only other country with similar information (New Zealand), the conservation of onychophorans seems to be of least concern for at least two thirds of the known species. *Epiperipatus isthmicola,* recently rediscovered after a century of absence in collections, can be considered Threatened because nearly all of its natural habitat has now been covered by a city.

## Abstract

**RESUMEN:** Las especies carismáticas, como el panda, desempeñan un papel importante en la conservación, y los gusanos de terciopelo posiblemente sean gusanos carismáticos. Gracias a su extraordinario mecanismo de caza, han inspirado a una banda de metal femenina en Japón, gusanos de origami en Rusia y monstruos de videojuegos en los Estados Unidos.

**Objetivo:** Evaluar su estado de conservación en Costa Rica.

**Métodos:** ubicamos todos los registros de recolección de las 29 especies conocidas de la Base de Datos Onychophora en el mapa de la Red de Conservación de Costa Rica.

**Resultados:** Encontramos que siete especies están protegidas dentro de reservas forestales, cinco en zonas protegidas, cuatro en refugios de vida silvestre, dos en parques nacionales y una, *Principapillatus hitoyensis,* en una reserva biológica estrictamente prístina. La especie más grande del mundo, *Peripatus solorzanoi,* se encuentra tanto dentro de una Reserva Forestal como en terrenos privados protegidos. La protección dentro de las áreas naturales de Costa Rica se aplica durante todo el año por personal que incluye guardias armados, y cuenta con el apoyo de programas educativos en las comunidades aledañas. Doce especies no se han encontrado en áreas protegidas, pero en Costa Rica, todas las especies biológicas, nombradas y no identificadas, están protegidas por la ley y no pueden ser legalmente recolectadas o exportadas, sin los permisos emitidos técnicamente.

**Conclusión:** Como en el único otro país con información similar (Nueva Zelanda), la conservación de los onicóforos parece ser menos preocupante para al menos dos tercios de las especies conocidas. Epiperipatus isthmicola, recientemente redescubierta después de un siglo de ausencia en colecciones, puede considerarse Amenazada porque casi toda su hábitat natural ha sido cubierto por una ciudad.

---

People are more likely to support the conservation of charismatic species like the panda, than of invertebrates like the cockroach (Courchamp et al., 2018). However, velvet worms can arguably be considered an exception: they are, to some extent, charismatic worms. They have inspired a female “death metal” band in Japan, origami worms in Russia and video game monsters in the USA, thanks to their extraordinary hunting mechanism (Monge-Nájera & Morera-Brenes, 2015).

Their conservation, though, is problematic because of contradictory and incomplete information. For decades, most authors have aligned with the idea that they are endangered because of their small populations and high susceptibility to habitat modification (e.g. Wells et al., 1983; Mesibov & Ruhberg, 1991; New, 1995; Vasconcellos et al., 2006). However, others have found that they survive forest fires (Mesibov & Ruhberg, 1991), volcanic eruptions (Barquero-González et al., 2016), deep habitat urbanization (Monge Nájera, 2018) and even the largest mass extinctions in the planet’s history (Monge-Nájera, 1995). The secret to their extraordinary survival is that, just like humans do at war, they hide underground until danger is over (Lavallard et al., 1975) and this has been a key factor in their evolution since the Cambrian (Monge-Nájera, 1995).

However, velvet worms should be taken into account in conservation programs because the tiny populations of cave species could easily disappear from overcollection (Peck, 1975, New, 1995) and because there are known cases of extinction apparently caused by habitat alteration. Three species, *Peripatopsis leonina, Peripatopsis clavigera* and *Opisthopatus roseus,* became extinct from habitat loss (Newlands & Ruhberg, 1978).

Two basic requirements to conserve a species are distinguishing it, and knowing where it occurs (Vasconcellos et al., 2006). Unfortunately, these two are difficult to meet for this phylum, because many species cannot be distinguished morphologically (Costa et al., 2018) and because they are not only rare, but also hard to find even in habitats where they occur (New, 1995). Without meeting these requirements, it is nearly impossible to know if their populations are endangered, and this is the key reason why they cannot be properly listed in conservation catalogues like the IUCN database (New, 1995).

Fully identifying species, their ranges, and their population status, is impossible at the time, but an assessment is still possible in some very particular cases. It has been attempted in New Zealand (Trewick et al., 2018), but the country is relatively large and has several onychophoran taxa that need revision (Oliveira et al., 2012). There is, nevertheless, one case in which conditions are more favorable: Costa Rica. The country is small and onychophorans have been actively collected for over a century; their geographic distribution is better known than anywhere else in the world (Barquero-González et al., 2016a). Additionally, even though many species have not been formally described, most have received common names and live specimens can be identified with photographic catalogues readily available to the public (Barquero-González et al., 2016b). Considering these favorable circumstances, we present here an assessment of the conservation status of all known Costa Rican species, based on their occurrence in the country’s comprehensive network of protected areas.

## MATERIALS AND METHODS

We tabulated all collection records from the Onychophora Database of the Escuela de Ciencias Biológicas, Universidad Nacional, Heredia, Costa Rica (Project 0094-17), and from the literature (Monge et al., 1993; Oliveira et al., 2012; Concha et al., 2014; Barquero-González et al., 2016a, 2016b, Giribet et al., 2018, Sosa-Bartuano et al., 2018). Additionally, we collected all available records from in-line sources as described in Barquero et al. (2016b), and from our own field observations, and we introduced them to the Costa Rica GIS of the Laboratorio de Ecología Urbana, Universidad Estatal a Distancia, San José (in full records Appendix 1). Protected areas are according to SINAC (Costa Rica, 2008).

## RESULTS

A total of 29 species have locality data allowing inclusion in the study; of these, 11 species only have known populations inside protected areas, most in Forest Reserves and Wildlife Reserves (four each); two in Protected Zones; one in a National Park and one in a Biological Reserve (Table 1). Six species have populations both inside and outside protected areas, mainly in Protected Zones and Forest Reserves (three each), and one inside a National Park (Table 2). Finally, twelve species do not have known populations inside protected areas (Table 2).

**TABLE 1.**
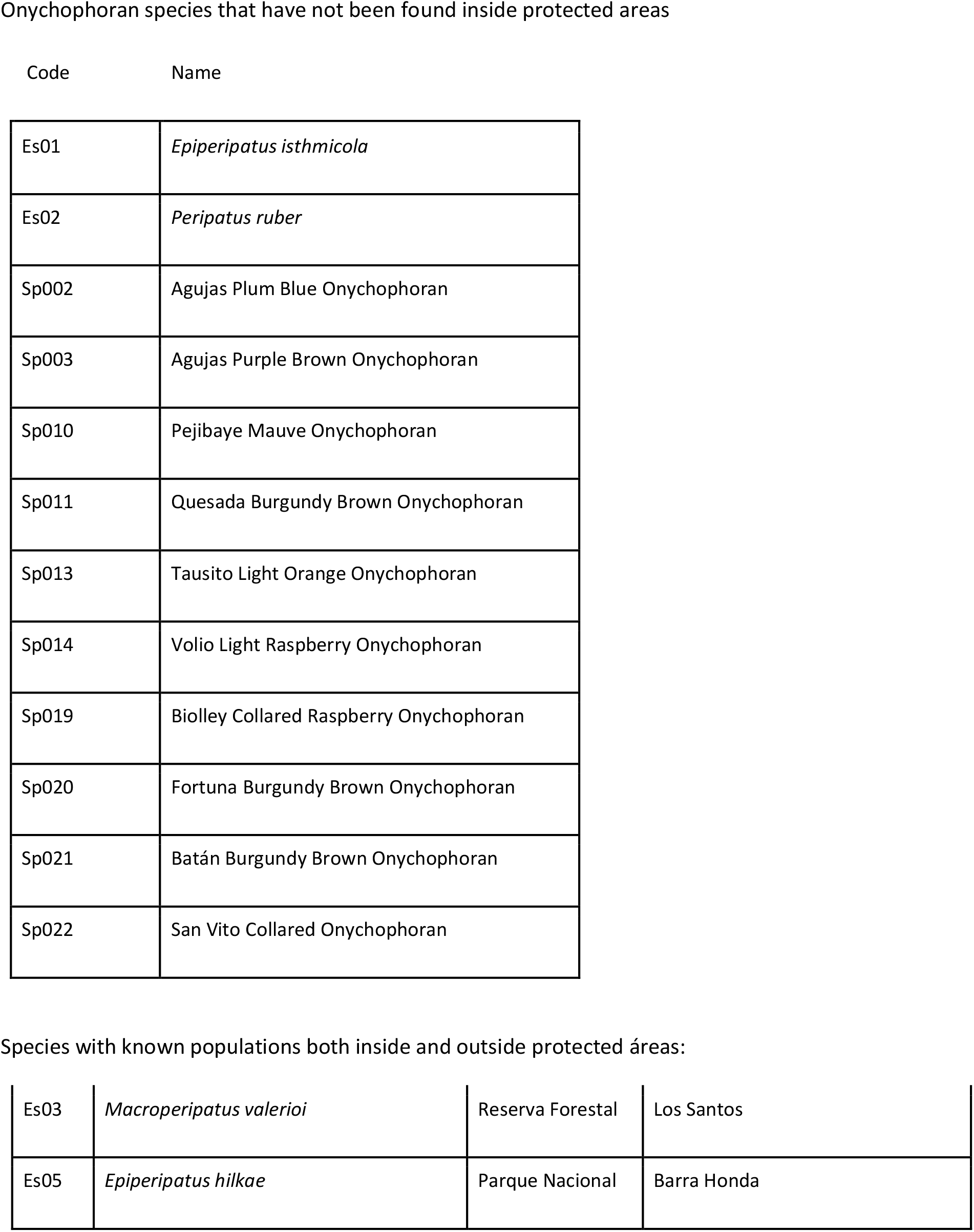
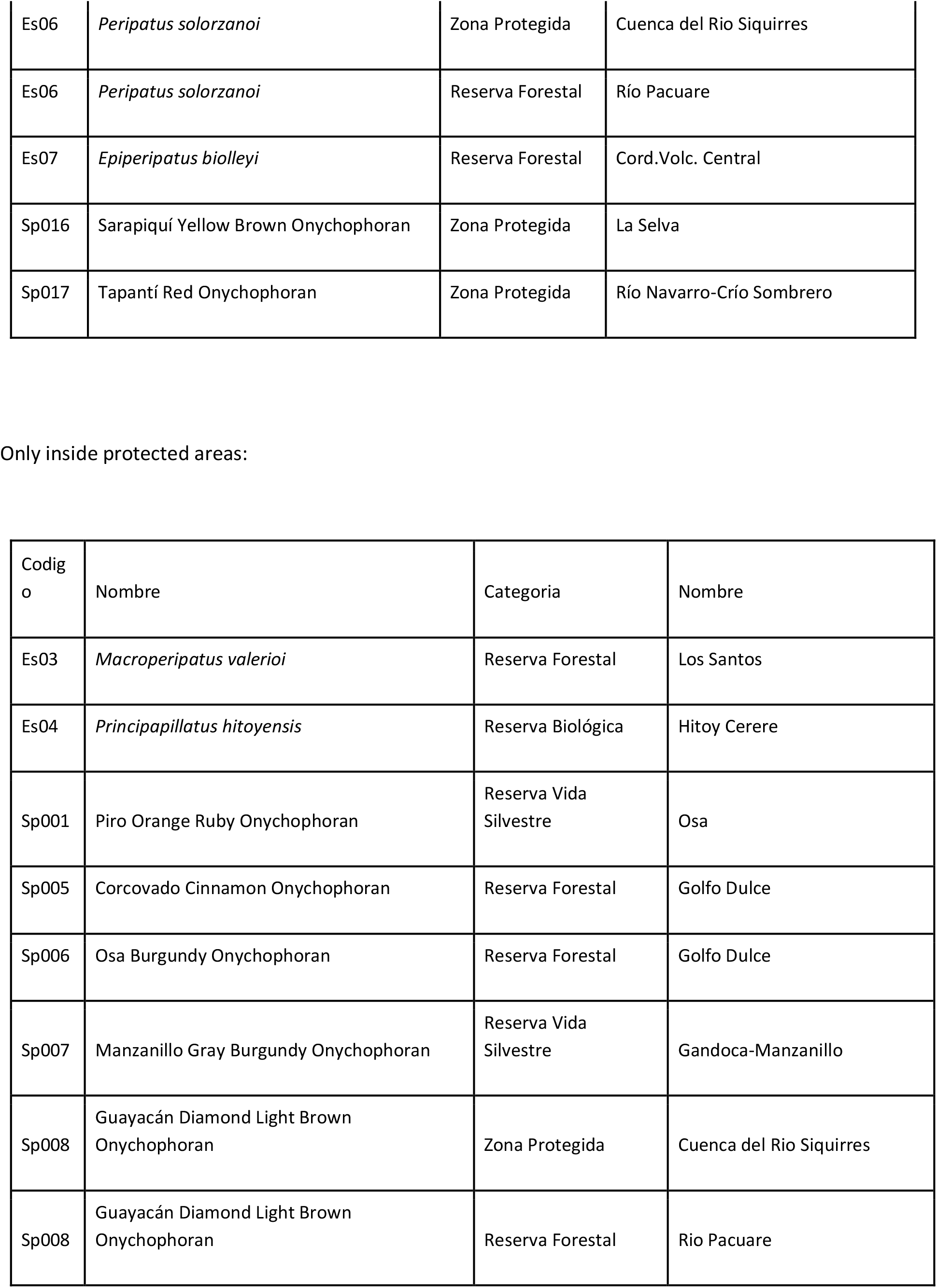
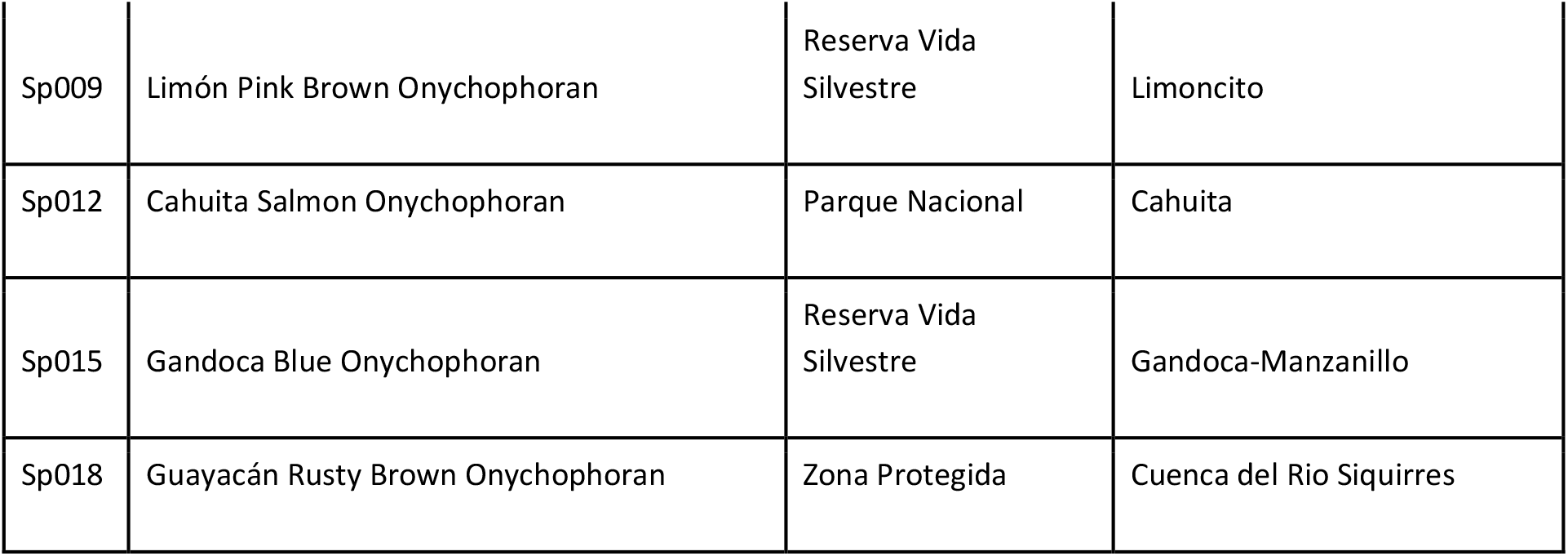
List of Costa Rican onychophoran species according to their distribution in relation with protected areas (inside, outside, or both).

| Code | Name |
| --- | --- |
| Es01 | *Epiperipatus isthmicola* |
| Es02 | *Peripatus ruber* |
| Sp002 | Agujas Plum Blue Onychophoran |
| Sp003 | Agujas Purple Brown Onychophoran |
| Sp010 | Pejibaye Mauve Onychophoran |
| Sp011 | Quesada Burgundy Brown Onychophoran |
| Sp013 | Tausito Light Orange Onychophoran |
| Sp014 | Volio Light Raspberry Onychophoran |
| Sp019 | Biolley Collared Raspberry Onychophoran |
| Sp020 | Fortuna Burgundy Brown Onychophoran |
| Sp021 | Batán Burgundy Brown Onychophoran |
| Sp022 | San Vito Collared Onychophoran |

Species with known populations both inside and outside protected áreas:
|  |  |  |  |
| --- | --- | --- | --- |
| Es03 | *Macroperipatus valerioi* | Reserva Forestal | Los Santos |
| Es05 | *Epiperipatus hilkae* | Parque Nacional | Barra Honda |
| Es06 | *Peripatus solorzanoi* | Zona Protegida | Cuenca del Río Siquirres |
| Es06 | *Peripatus solorzanoi* | Reserva Forestal | Río Pacuare |
| Es07 | *Epiperipatus biolleyi* | Reserva Forestal | Cord.Volc. Central |
| Sp016 | Sarapiquí Yellow Brown Onychophoran | Zona Protegida | La Selva |
| Sp017 | Tapantí Red Onychophoran | Zona Protegida | Río Navarro-Crío Sombrero |

Only inside protected areas:
| Codig<br>o | Nombre | Categoría | Nombre |
| --- | --- | --- | --- |
| Es03 | *Macroperipatus valerioi* | Reserva Forestal | Los Santos |
| Es04 | *Principapillatus hitoyensis* | Reserva Biológica | Hitoy Cerere |
| Sp001 | Piro Orange Ruby Onychophoran | Reserva Vida Silvestre | Osa |
| Sp005 | Corcovado Cinnamon Onychophoran | Reserva Forestal | Golfo Dulce |
| Sp006 | Osa Burgundy Onychophoran | Reserva Forestal | Golfo Dulce |
| Sp007 | Manzanillo Gray Burgundy Onychophoran | Reserva Vida Silvestre | Gandoca-Manzanillo |
| Sp008 | Guayacán Diamond Light Brown Onychophoran | Zona Protegida | Cuenca del Río Siquirres |
| Sp008 | Guayacán Diamond Light Brown Onychophoran | Reserva Forestal | Río Pacuare |
| Sp009 | Limón Pink Brown Onychophoran | Reserva Vida<br>Silvestre | Limoncito |
| Sp012 | Cahuita Salmon Onychophoran | Parque Nacional | Cahuita |
| Sp015 | Gandoca Blue Onychophoran | Reserva Vida<br>Silvestre | Gandoca-Manzanillo |
| Sp018 | Guayacán Rusty Brown Onychophoran | Zona Protegida | Cuenca del Río Siquirres |

By kind of protection, seven species are protected inside forest reserves, where only some sustainable use of the forest is allowed, requiring conservation of the soil and vegetation where onychophorans and their prey live. Five species are inside Protected Zones, where human activity is allowed as long as soil, water and vegetation are not damaged. Four are in Wildlife Refuges, where some private activities and even human settlements are allowed, but always under sustainable management practices. Two occur inside National Parks, where nature is kept unaltered except for a small area where research, education and ecotourism are allowed. Finally, one species, Principapillatus hitoyensis, is protected inside a Biological Reserve, where the goal is to conserve a pristine natural state. It must be borne in mind that protection inside Costa Rican nature areas is enforced year round by trained personnel that includes armed guards, and supported by educational programs in surrounding communities (http://www.sinac.go.cr/EN-US/Pages/default.aspx).

## DISCUSSION

Since the 19th century, most authors mentioned the rarity of onychophorans, but their conservation literature may have started 30 years ago with the pioneer work of Robert Mesibov in Tasmania (Mesibov & Ruhberg, 1991). Scattered articles followed (New, 1995; Gleeson, 1996; Vasconcellos et al., 2006; Daniels, 2011), and the subject of their survival in cities is even more recent and began simultaneously in Costa Rica and New Zealand (Barquero-González, et al., 2016; Barret et al., 2016; Monge-Nájera 2017, 2018).

Currently, while there are no conservation studies for most of the nearly 200 named species of velvet worms, the IUCN considers four in critical danger (in South Africa and Brazil), two endangered (in Tasmania and Jamaica), four vulnerable (in South Africa and New Zealand) and one in low risk (in Jamaica), mainly from habitat loss (Oliveira et al., 2015; IUCN 2018). The Estacão Ecológica do Tripuí in Minas Gerais, Brazil, was created in part to protect *Peripatus acacioi* (http://www.wikiparques.org) and efforts to conserve an undescribed urban species were also done in Dunedin, New Zealand, with a strong citizen participation (Monge-Nájera & Morera-Brenes, 2015; Barrett et al., 2016).

Four species deserve individual consideration, *Epiperipatus biolleyi, Principapillatus hitoyensis, Peripatus solorzanoi,* and *E. isthmicola.* Biolley’s onychophoran, *E. biolleyi,* has been collected both inside the cloud forest of Braulio Carrillo National Park, and in cattle farms near the Irazú volcano, where it survived the large eruption of 1963-1965; it is among the best known species in the world by the number and depth of studies that have been published about it (Barquero-González et al., 2016). Considering this record, the survival of *E. biolleyi* is probable in the foreseeable future.

The only known member of the genus *Principapillatus,* the Caribbean species *P. hitoyensis,* may be the Costa Rican species with the best taxonomic data, because it was described recently with both DNA information and morphological detail that exceeds previous descriptions (Oliveira et al., 2012). This species from lowland rainforest is known only from the Hitoy Cerere Biological Reserve but can also live among the roots of banana trees, suggesting that it could expand its range into nearby banana plantations. In any case, the protection level in the reserve makes its survival likely. The importance of *P. solorzanoi* cannot be overstated because it is the largest onychophoran in the world (Morera-Brenes & Monge-Nájera, 2010). Luckily, it occurs both inside the Río Pacuare Forest Reserve, and in private land where, at least at the time of this study, it is also protected by the owners.

Finally, *E. isthmicola* is extraordinary because it was thought to be extinct for almost a century, after its description from what was originally tropical Premontane Moist Forest, then pasture land, and finally the heavily urbanized downtown of San José city in central Costa Rica. This species currently is known only from the original description and from a few “recent” collections inside the city core (Barquero-González, et al., 2016). The fact that it can hardly be protected in the middle of a city is concerning, and makes it a particularly fit species for a conservation campaign.

A recent analysis of 12 onychophoran species in New Zealand reported eight Not Threatened, with large, stable populations; three At Risk because they are naturally uncommon; and one Data Deficient (none were Threatened with extinction, Trewick et al., 2018). If that classification is applied to the Costa Rican species, all can be considered At Risk because all are naturally uncommon, and because in our experience, they are harder to find now than 30 years ago in places like Coronado, the habitat of *E. biolleyi.* A few years ago, the Data Deficient category, which includes species that may be extinct but lack proper data, would have included *E. isthmicola,* only rediscovered in 2004 (Barquero-González et al., 2016a). It could be considered, however, Threatened (an equivalent of IUCN’s Endangered), because nearly all of its natural habitat has now been covered with concrete (Barquero-González et al., 2016a).

At the time of this report, there are only two countries with conservation assessments for “all” of their onychophoran species; New Zealand and Costa Rica. Nevertheless, New Zealand is five times the size of Costa Rica (which has an estimated of more than 50 species, Morera-Brenes and Barquero-González: unpublished); by area alone, New Zealand may have around 250 species, and the 2018 conservation assessment based on 12 species might under represent the country’s onychofauna.

In conclusion, our results show that about two thirds of the known Costa Rican species are protected inside properly enforced conservation areas, and the rest (12 species known to date) occur in unprotected private property, including cities. However, this does not mean that they fully lack protection, because in Costa Rica, all biological species, named or unnamed, are protected by law and cannot be legally collected, or exported, without technically issued permits (MAG 2008). Overprotection, which takes place when bureaucrats become an unreasonable barrier to research, can also be deleterious to onychophoran conservation (New, 1995), but --in our experience--this is not currently a problem for research on Costa Rican onychophorans.

The conservation of their naturally low (but catastrophe-resistant?) populations appears to be of least concern for at least two thirds of the known onychofauna of Costa Rica

**Figure 1.**
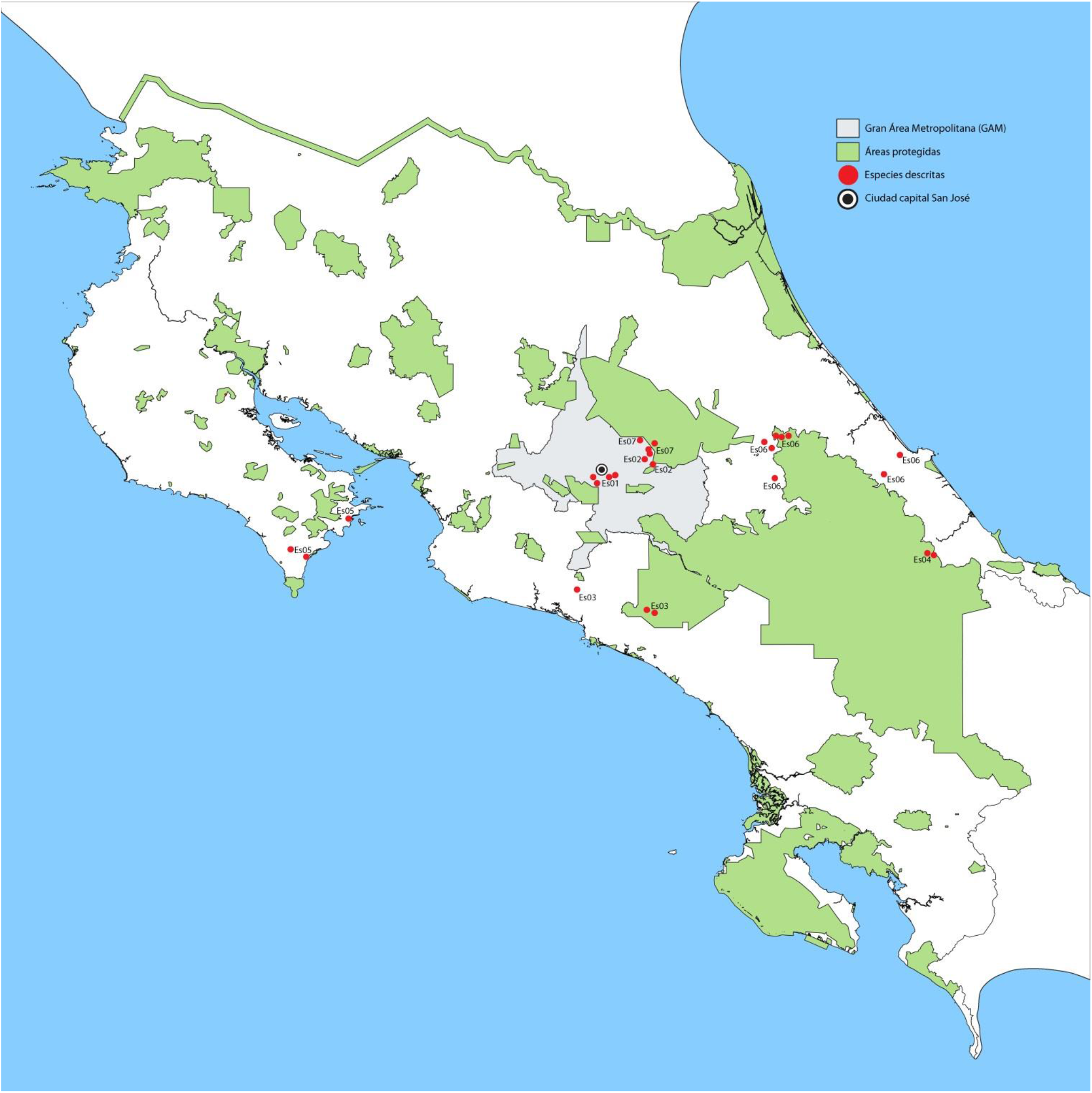
Collection localities of formally described Costa Rican onychophoran species in relation with protected areas. Species codes in Table 1.

**Figure 2.**
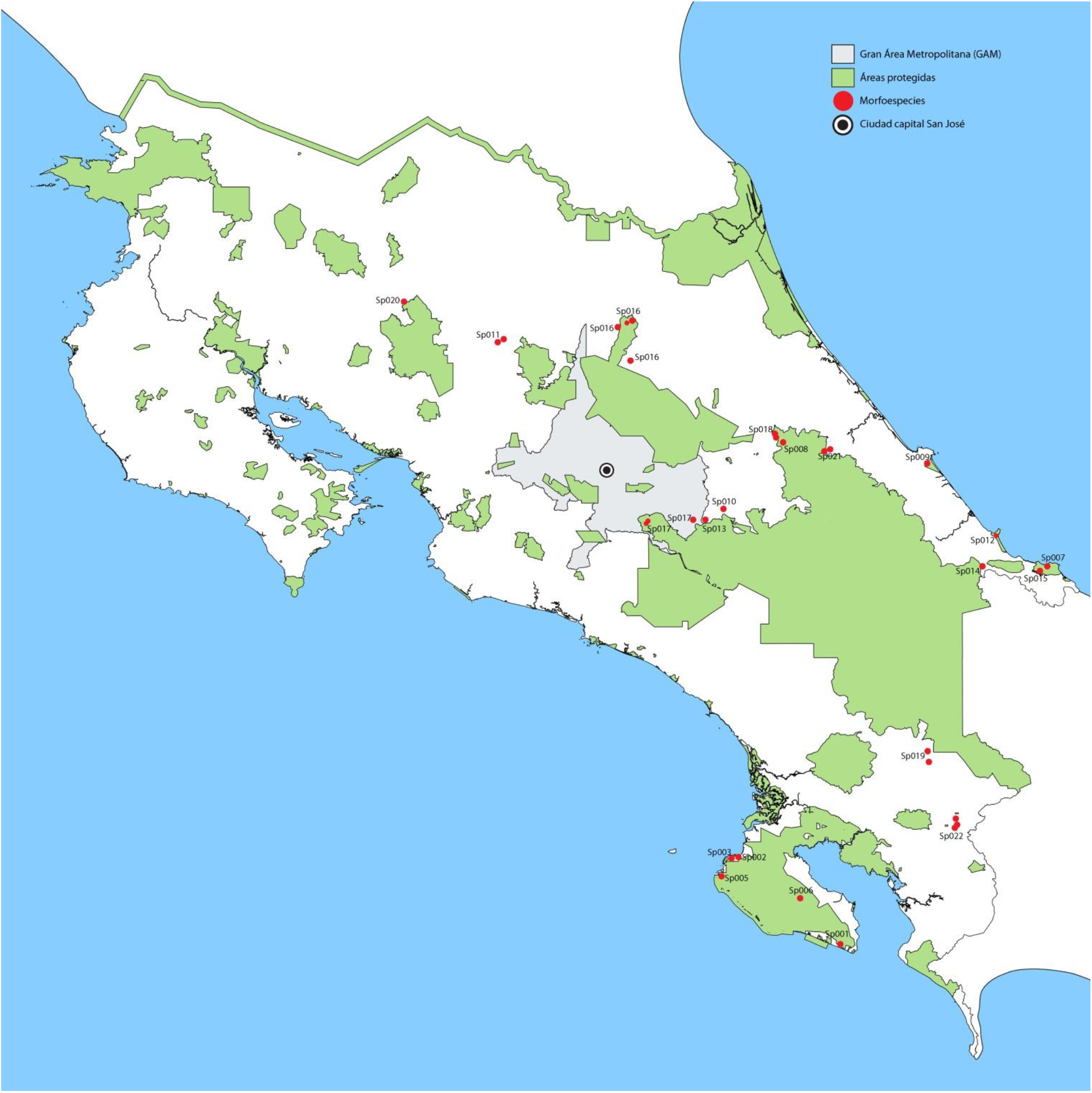
Collection localities of unnamed Costa Rican onychophoran species in relation with protected areas. Species codes in Table 1.

## ACKNOWLEDGMENTS

We thank Zaidett Barrientos for her support and cooperation, Carolina Seas and Maribel Zúñiga for their assistance and XXX for comments to improve an earlier draft. This study was partially financed by Project UNA 0094-17, Universidad Nacional de Costa Rica and done with the support of Laboratorio de Ecología Urbana, UNED.

## APPENDIX 1 DETAILED COLLECTING LOCALITIES

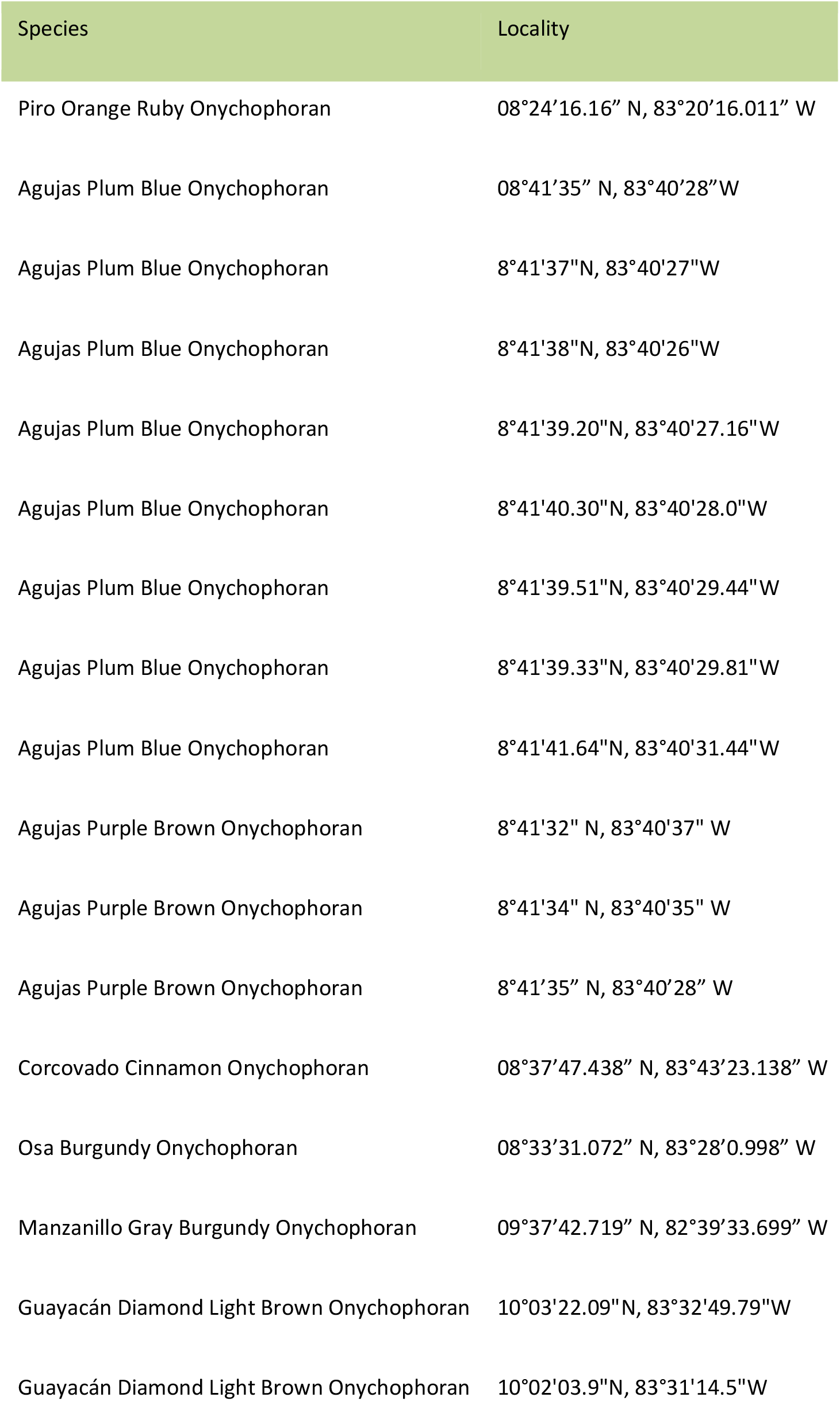
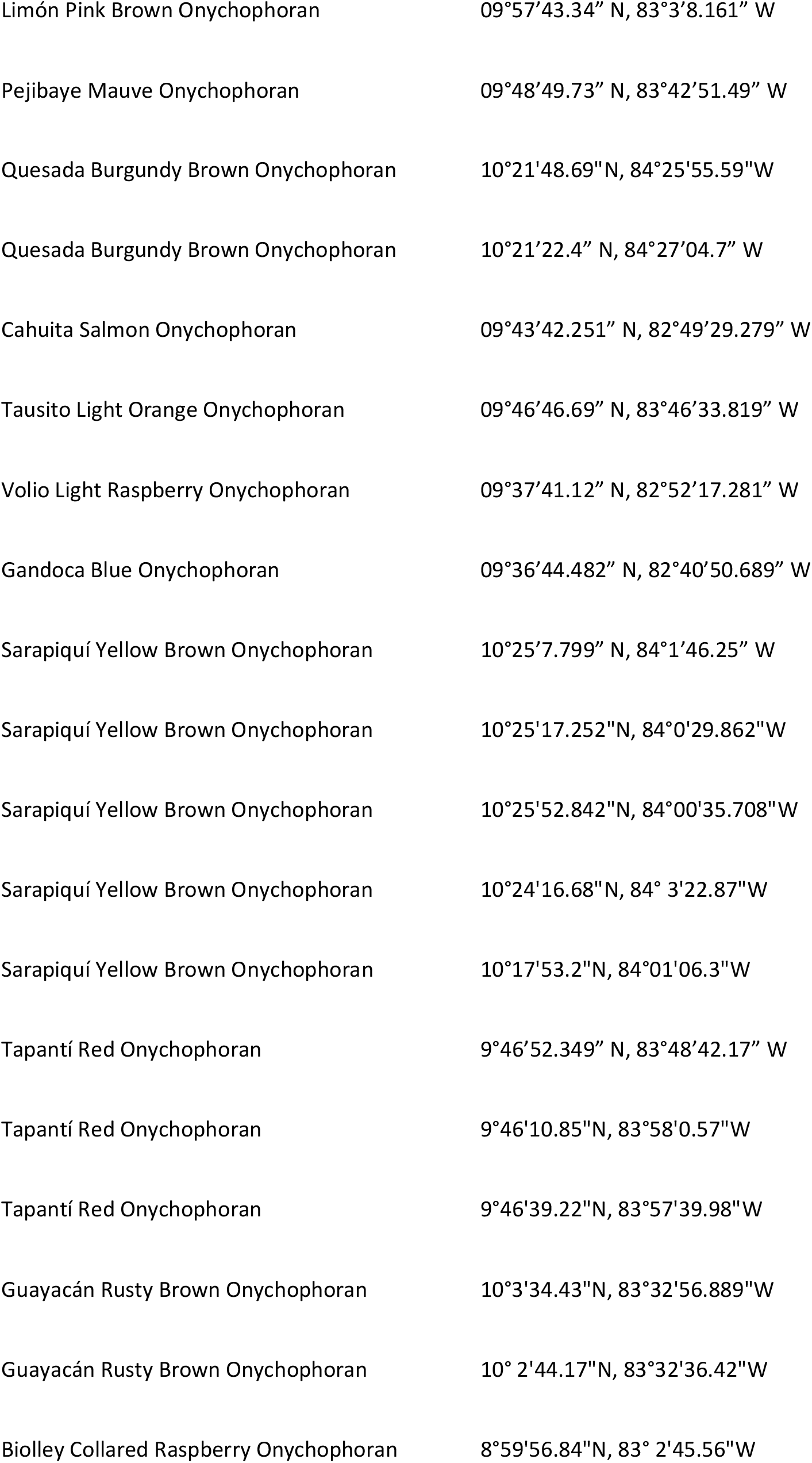
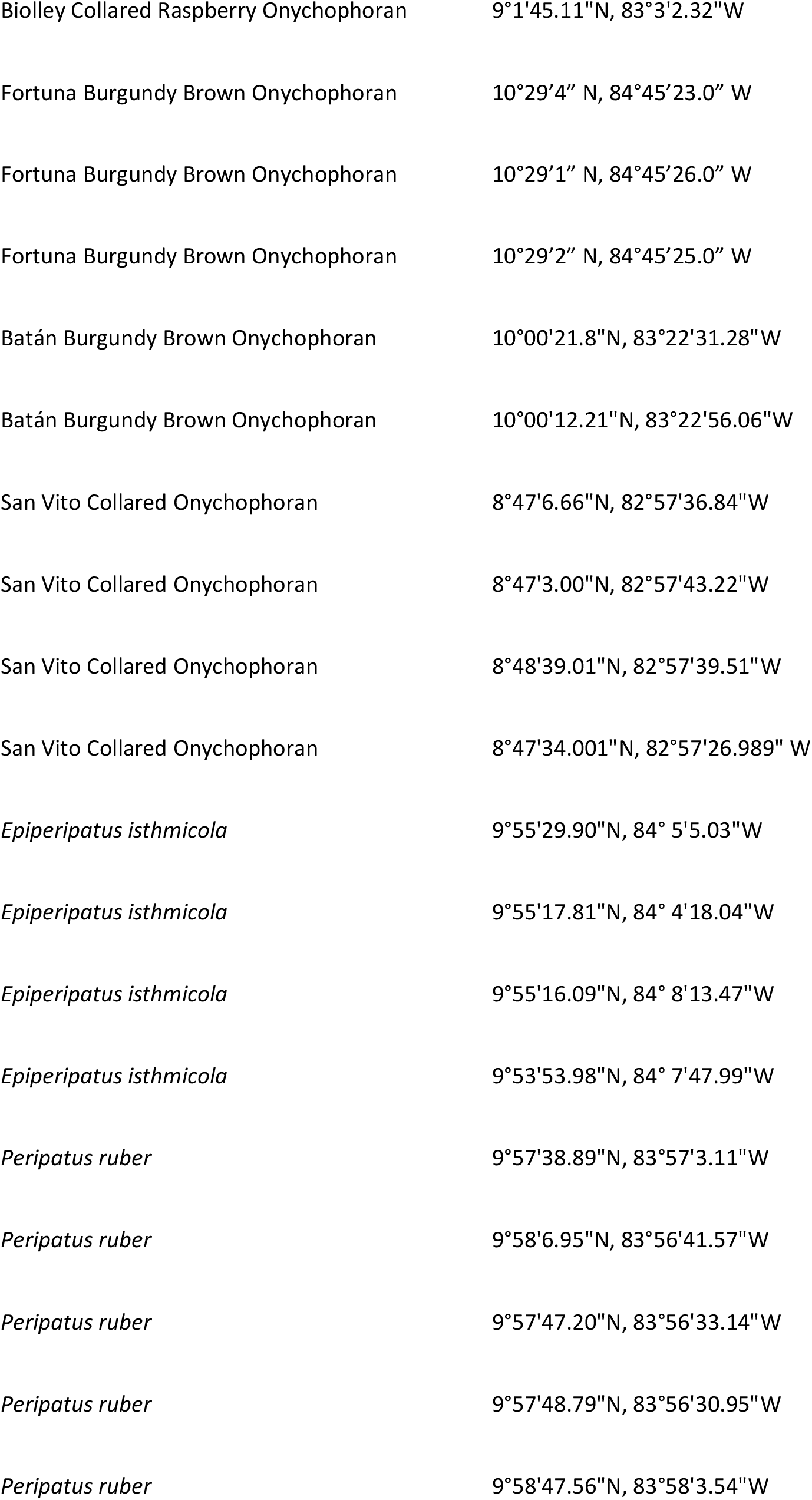
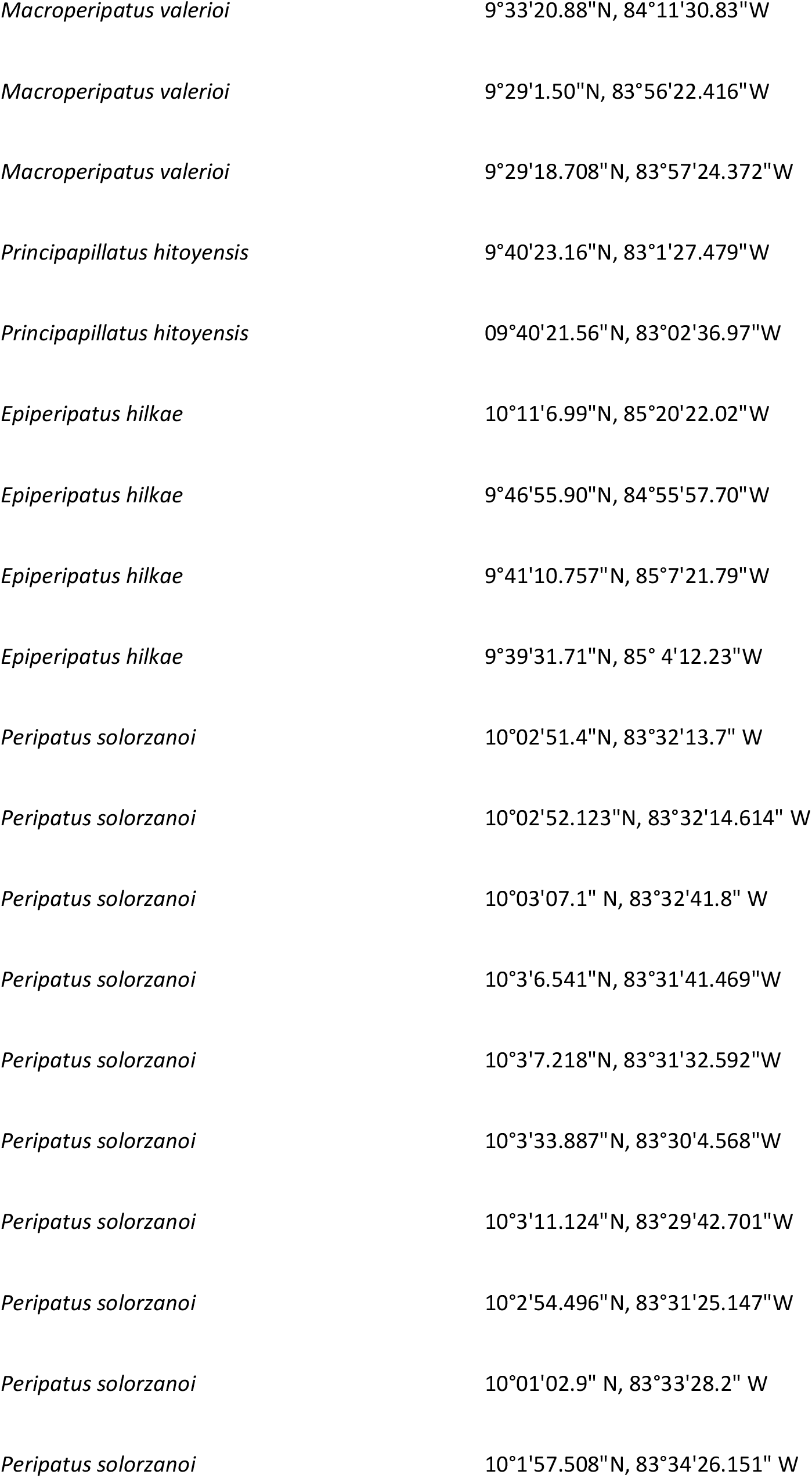
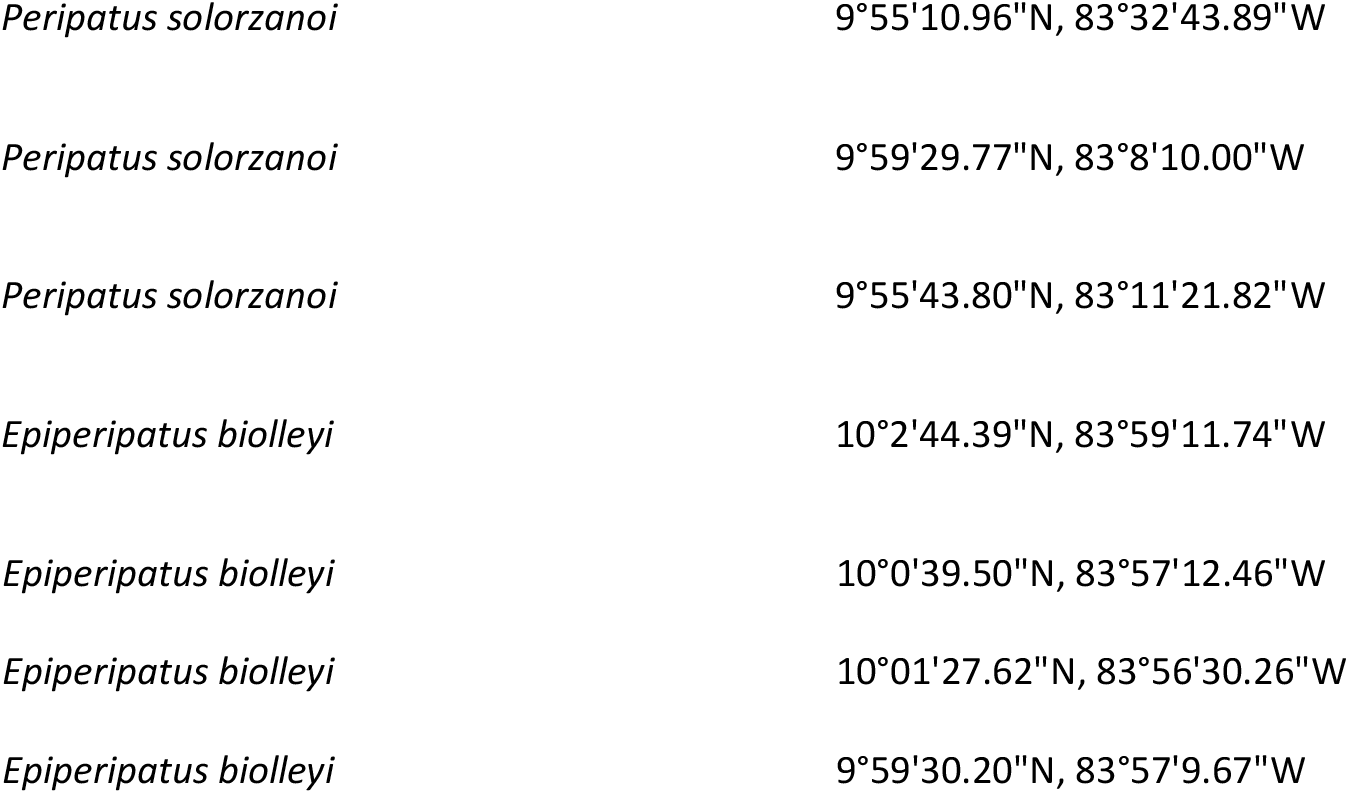

